# Discovery of a non-canonical GRHL1 binding site using deep convolutional and recurrent neural networks

**DOI:** 10.1101/2022.06.28.497553

**Authors:** Sebastian Proft, Janna Leiz, Udo Heinemann, Dominik Seelow, Kai M. Schmidt-Ott, Maria Rutkiewicz

## Abstract

Transcription factors regulate gene expression by binding to transcription factor binding sites (TFBSs). Most models for predicting TFBSs are based on position weight matrices (PWMs), which require a specific motif to be present in the DNA sequence and do not consider interdependencies of nucleotides. Novel approaches such as Transcription Factor Flexible Models or recurrent neural networks consequently provide higher accuracies. However, it is unclear whether such approaches can uncover novel non-canonical, hitherto unexpected TFBSs relevant to human transcriptional regulation. In this study, we trained a convolutional recurrent neural network with HT-SELEX data for GRHL1 binding and applied it to a set of GRHL1 binding sites obtained from ChIP-Seq experiments from human cells. We identified 46 non-canonical GRHL1 binding sites, which were not found by a conventional PWM approach. Unexpectedly, some of the newly predicted binding sequences lacked the CNNG core motif, so far considered obligatory for GRHL1 binding. Using isothermal titration calorimetry, we experimentally confirmed binding between the GRHL1-DNA binding domain and predicted GRHL1 binding sites, including a non-canonical GRHL1 binding site. Mutagenesis of individual nucleotides revealed a correlation between predicted binding strength and experimentally validated binding affinity across representative sequences. This correlation was neither observed with a PWM-based nor another deep learning approach. Our results show that convolutional recurrent neural networks may uncover unanticipated binding sites and facilitate quantitative transcription factor binding predictions.

## INTRODUCTION

Transcription factors are proteins that regulate gene expression by binding to specific DNA sequences, referred to as transcription factor binding sites (TFBSs). Traditionally, TFBSs have been characterised by positional weight matrices (PWMs), providing a probabilistic representation of binding motifs (1,2). Today, PWMs are derived from DNA sequences obtained from high-throughput experiments such as high-throughput systematic evolution of ligands by exponential enrichment (HT-SELEX) or chromatin immunoprecipitation with sequencing (ChIP-Seq) (3-5). PWMs have been determined for various transcription factors across multiple species (6,7). However, PWMs have limitations, including the assumed independence of nucleotides within the sequence and a poor representation of motifs with low binding affinities (8). Transcription factor flexible models (TFFMs) are based on Hidden Markov Models and overcome some of these limitations by conserving first-order interactions between the bases (9).

The significant advances in deep learning technologies over the last decade have also led to the development of artificial neural networks (ANNs), such as convolutional and recurrent neural networks, for the identification of TFBSs (for recent reviews, see (10-13)). ANNs extract binding motif characteristics directly from sequencing data, learn higher-level interdependencies between all bases, and encompass different forms of architectures. While the training data must be pre-labelled (i.e. binding vs. non-binding), learning of dependencies between the nucleotides occurs automatically during the training process. For example, DeepBind, a method published in 2015, uses deep convolutional neural networks (CNNs) to discover binding patterns of transcription factors in experimental data (14). DeepBind highlights the potential of ANNs to find new binding sites and to predict the effect of DNA variants on transcription factor binding. More recent models also incorporate recurrent neural networks (RNNs) (15-17), which are better suited to process sequential data (18).

The transcription factor grainyhead-like 1 (*GRHL1* gene) belongs to the family of Grh transcription factors first described in drosophila (19,20). *GRHL1* is mainly expressed in the developing surface ectoderm and hair follicles as well as the developing and mature epidermis (21). It plays a critical role in the formation of desmosomes between keratin-expressing epithelial cells and directly regulates the expression of desmoglein 1 (*DSG1*) (22). Consequently, *Grhl1*-null mice exhibit abnormal hair and skin phenotypes, such as delayed coat growth, premature hair loss, and palmoplantar keratoderma (22), a congenital disorder also found in humans (23). Additionally, several studies implicate a functional role for the GRHL1 protein in cancer (24-26), further underscoring its potential role in genetic disease. The highly conserved GRHL1 consensus motif (5’-AACCGGTT-3’) has long been known (27) and the structural basis of its recognition by the GRHL1 DNA-binding domain has been unveiled (28). So far, no additional binding motifs have been reported. However, it is crucial to identify all potential binding sites throughout the genome to fully understand the underlying molecular basis of transcriptional regulation, regulatory networks, genomic traits, and genetic disease (29-32) mediated by GRHL1.

In this work, we trained, validated, and tested a neural network that combines the technologies of CNN and RNN on HT-SELEX data enriched for GRHL1. For simplicity, our model will be referred to as RNN. We next tested the ability of the RNN to discriminate true GRHL1 binding sites from binding sites for another transcription factor (MYOD) in human ChIP-Seq data. We compared the performance of the RNN with the classical PWM-based approach using Find Individual Motif Occurrences (FIMO (33)), and with DeepBind (14). The RNN accurately identified binding sites with the long-known GRHL1 consensus binding motif but also discovered novel non-canonical GRHL1 binding sites that were missed entirely by FIMO’s traditional PWM-based probabilistic method as well as by DeepBind. In addition, we tested whether the RNN could predict the quantitative effect of single nucleotide variants on GRHL1 binding affinity. GRHL1 binding affinities to a spectrum of representative DNA sequences, including non-canonical binding sites, were validated experimentally using isothermal titration calorimetry (ITC) experiments.

## MATERIAL AND METHODS

### GRHL1 expression and purification

The GRHL1-DNA binding domain construct (UniProtKB: Q9NZI5, aa 248 – 485) was cloned into the pQlinkH vector (34) as reported earlier (28). It was transformed into *Escherichia coli* BL 21 (DE3) – T1R chemocompetent cells. Large-scale cultures were induced at 18 °C with 0.5 mM isopropyl β-D-1-thiogalactopyranoside (IPTG). The harvested cells were resuspended in lysis buffer (1 x PBS pH 7.5, 500 mM NaCl, 5% glycerol, 0.5 mM DTT, 40 mM imidazole, 1 mg/ml lysozyme, 1 mg/ml DNAse supplemented with 1 Pierce protease inhibitor tablet (Pierce, USA) per 50 ml of buffer). The supernatant was applied on a HisTrapTM FF Crude (Cytiva, USA) column equilibrated with H4 buffer (1 x PBS pH 7.5, 500 mM NaCl, 5% glycerol, 0.5 mM DTT, 40 mM imidazole) and eluted with H10 buffer (1 x PBS pH 7.5, 500 mM NaCl, 5% glycerol, 0.5 mM DTT, 400 mM imidazole). The eluent was dialysed vs. H4 buffer, and the His6-tag was cleaved off with TEV protease. The tag was bound to HisTrapTM FF Crude, at the same time the protein-containing flow-through fractions were concentrated and applied onto HiLoad 16/600 Superdex 200 pg columns (Cytiva, USA) equilibrated with ITC buffer (20 mM HEPES-NaOH pH 7.2, 125 mM NaCl, 2 mM DTT). The sample was concentrated using Vivaspin filters to up to 156 μM for the first batch and to 200 μM for the second. All purification steps were performed at 4 °C. The protein purification was controlled through SDS-PAGE electrophoresis.

### Double-stranded (ds) DNA preparation

Selected single-stranded DNAs and their reverse complementary fragments were synthesised by Eurofins, Luxemburg. Samples were dissolved in nuclease-free water (Thermo Fisher Scientific, USA). Their concentration was measured in triplicates using NanoDrop OneC (Thermo Fisher Scientific, USA). The forward and reverse fragments were mixed in a 1: 1 molar ratio and heated up to 60 °C. The samples were slowly (1 °C/5 min) cooled down to room temperature and then incubated at 10 °C for another two hours. dsDNA samples were dialysed at 10 °C overnight vs. ITC buffer and diluted to a concentration of approximately 10 μM.

### Isothermal titration calorimetry (ITC) experiments

We used a MicroCal PEAQ-ITC microcalorimeter (Malvern Panalytical GmbH, Germany) for ITC measurements. Experiments were performed in ITC buffer (20 mM HEPES-NaOH pH 7.2, 125 mM NaCl, 2 mM DTT). GRHL1 was titrated in 19 or 24 steps (1.5-2 μl) into a dsDNA ligand solution in a calorimeter cell. The reference cell contained the ITC buffer. The mean signal resulting from titrating GRHL1 into ITC buffer was subtracted. After initial testing, the dsDNA:protein ratio was fixed as 1:2 to reflect the known binding model of two GRHL1 DNA-binding domains binding one motif-containing dsDNA molecule. Measurements were performed in triplicate for each ligand. Data analysis (baseline adjustment, peak integration, normalisation of the reaction heats, data fitting, and evaluation) was performed using the MicroCal PEAQ-ITC Analysis software.

## RESULTS

### Training, validation, and selection of the RNN model on in vitro data

We used the same GRHL1 binding sequences as employed by DeepBind from a HT-SELEX experiment, consisting of 203,209 artificially created sequences of 20 nucleotides (5) to build our RNN model. To generate control, non-binding sequences, we shuffled the position of each base, thereby breaking up the binding site patterns while keeping the abundance of each dinucleotide and overall GC content consistent (35). This shuffle was achieved using “fasta-shuffle-letters” with k-mer set to 2 from the meme suite version 5.3.3 (36). The dataset was then randomly split into training, validation, and testing subsets in the ratio of 70% / 15% / 15%. This process created a balanced dataset with an equal abundance of binding and non-binding sequences.

We built several RNNs in python using the pysster package (37) on the HT-SELEX training data subset using various hyperparameters (**Figure 1A**). Hyperparameters, such as the model architecture or the dropout ratios, control the learning process and are essential for optimising the model. They cannot be derived from the data in the training process and must therefore be chosen beforehand. Selecting the best hyperparameters can be done by comparing the performance of the different models after training has concluded.

**Figure 1.**
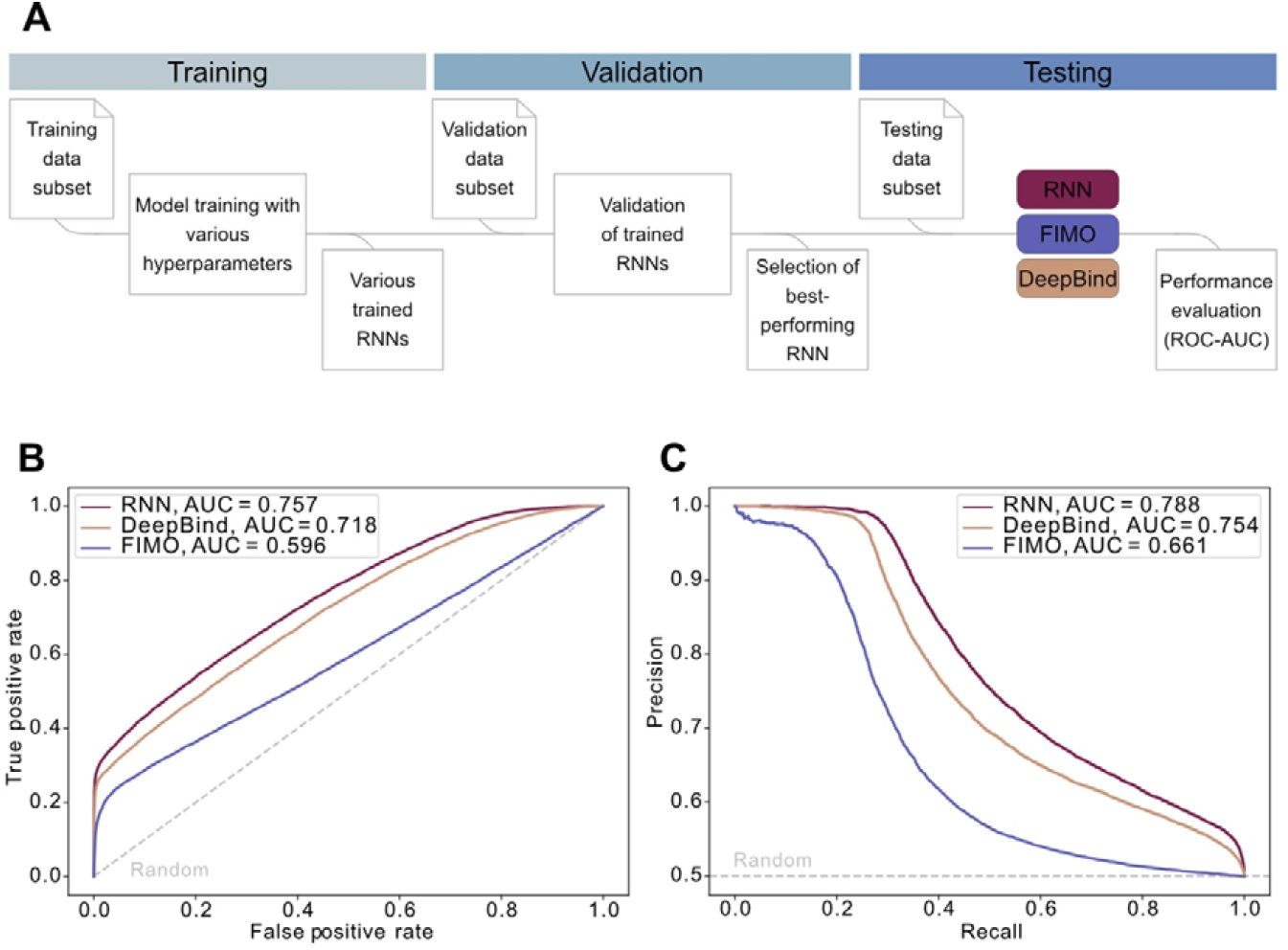
Performance of the RNN model on HT-SELEX data. (A) Workflow for training, validation, and testing of the RNN model on HT-SELEX data. The dataset (203,209 sequences) was split into a training, validation, and testing subset in the ratio 70% / 15% / 15%. The final performance in the testing data subset was evaluated for the selected RNN model, FIMO, and DeepBind. (B) Area under the curve (AUC) for the receiver operator characteristic (ROC) curve of the tested models. (C) AUC for the receiver precision-recall curve of the tested models. The grey dotted line depicts random predictions with an AUC of 0.5.

Here, we trained RNNs with and without a convolutional layer, with different lengths for the kernel in the convolutional layer, different dropout ratios for the dense layer, and with a uni- or bidirectional RNN architecture (**Supplementary Table 1**). Final hyperparameters (**Supplementary Table 1**) were selected based on their performance as indicated by the area under the curve (AUC) of the receiver operator characteristic (ROC) curve on the validation data subset (**Figure 1A**).

To ensure that our selected model performs well, we compared it against the established probabilistic approach of FIMO (applying the PWM MA0647.1 from JASPAR (7)) and the GRHL1 DeepBind model (14), by applying all three models to the same previously unseen test dataset (**Figure 1A**). The final performance was assessed by calculating the AUC on ROC (**Figure 1B**) and precision-recall curves for each model (**Figure 1C**). Our final RNN (architecture depicted in **Figure 2**) outperformed FIMO and was on par with DeepBind on the HT-SELEX test dataset.

**Figure 2.**
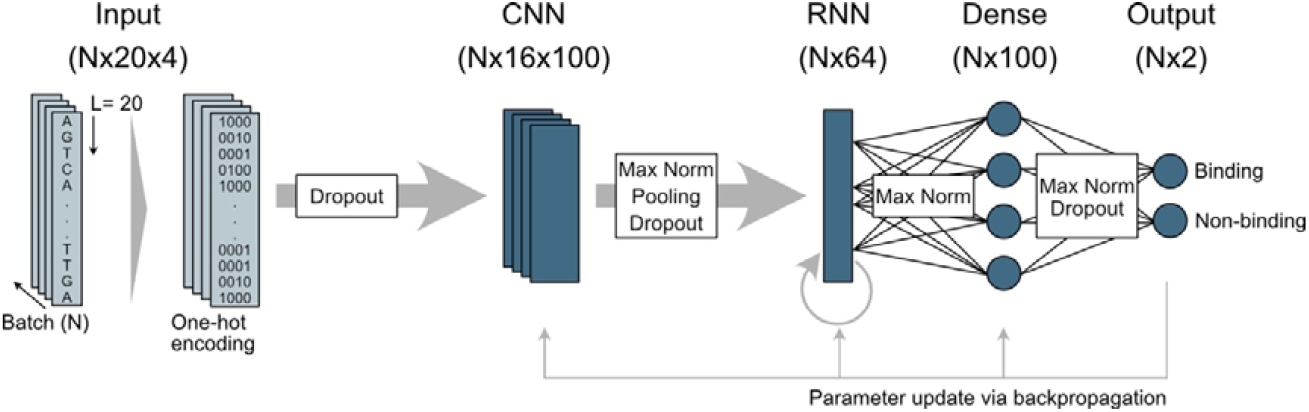
Architecture of the final RNN model. Input sequences (N sequences of length 20) were transformed into numerical data using one-hot encoding. The model used a convolutional layer (CNN) and a recurrent layer (RNN) to extract features from the input sequences. Model regularization techniques like dropout, max pooling, and max norms were used throughout the model to prevent overfitting. The extracted features were fed into a dense, or fully connected layer. As a result, sequences were classified as “binding” or “non-binding” by the final dense layer (output layer). Model optimisation was achieved by parameter update via backpropagation.

### The RNN model performs better than other approaches on in vivo data

After successful model selection, we ran the RNN, FIMO, and DeepBind on 7,857 high-confidence GRHL1-DNA binding sites within the human genome from a ChIP-Seq experiment in human MCF-7 cells (SRA accession number SRX7122002) to test their performance on never-before-seen in vivo data (38). As a negative dataset we used 9,509 MYOD binding sequences from a ChIP-Seq experiment in myoblasts (SRA accession number SRX341010) (39). MyoD had the least overlap with GRHL1 binding sites of the negative datasets tested. We used the intersect function from BEDtools (version 2.30.0) (40) to confirm minimal overlap of genomic regions between the positive and negative datasets. GRHL1 and MYOD ChIP-Seq peaks showed only 201 overlapping sequences that were removed from the negative data set.

We used sequences of 100 nucleotides centred around the ChIP-Seq peak summit (qval < 1E-05 for GRHL1 and qval < 1E-20 for MYOD). As the RNN was trained to score sequences of 20 nucleotides, we used a sliding-window approach to identify the highest-scoring 20-mer within each ChIP-Seq peak. Score ranges vary widely between methods, with high scores within each range indicating a sequence is designated as binding, while low scores imply no binding. In the case of our RNN, scores can range from 0 to 1, with scores > 0.5 denoting a positive binding prediction. The RNN model showed a clearer difference in score distribution between the positive GRHL1 and negative MYOD class than both FIMO and DeepBind (**Figure 3**). Individual prediction scores and respective percentiles for each GRHL1 and MYOD sequence can be found in **Supplementary Tables 2** and **3**.

**Figure 3.**
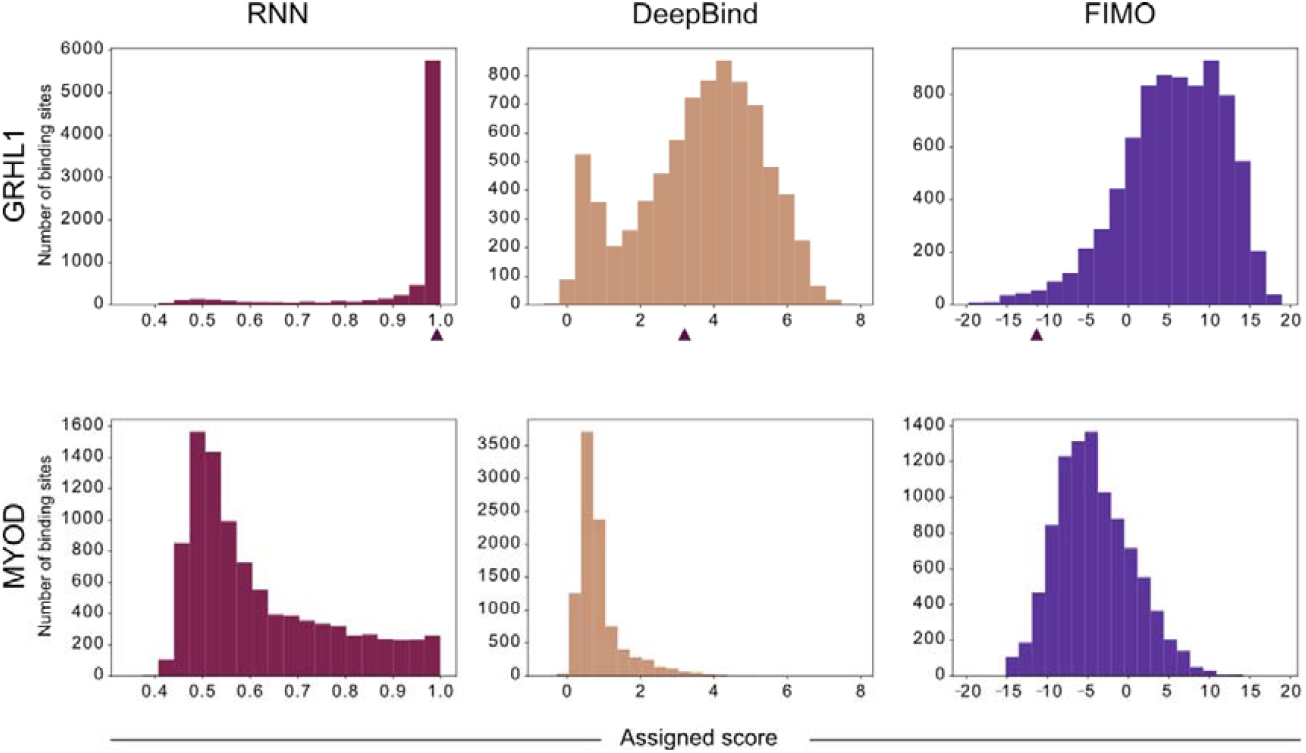
Score distribution assigned by the three models for GRHL1 and MYOD binding sites. Note that the different approaches have different score ranges, with higher values representing stronger binding within each approach. The arrowheads indicate the scores assigned to a GRHL1 ChIP-Seq peak containing a novel non-canonical GRHL1 sequence identified by the RNN model (“Novel”, see below).

To compare the RNN, FIMO, and DeepBind, we calculated ROC curves for each model on 16 bins of 500 GRHL1 ChIP-Seq peaks, sorted by decreasing ChIP-Seq score for GRHL1 binding (**Figure 4A**). As the negative dataset we picked a random set of 500 MYOD binding sites for each bin. The calculation was repeated 1000 times with different randomly picked MYOD sequences to reduce potential effects of the selection of non-binding sequences on the prediction (randomly selected peaks for each run see **Supplementary Table 4**).

**Figure 4.**
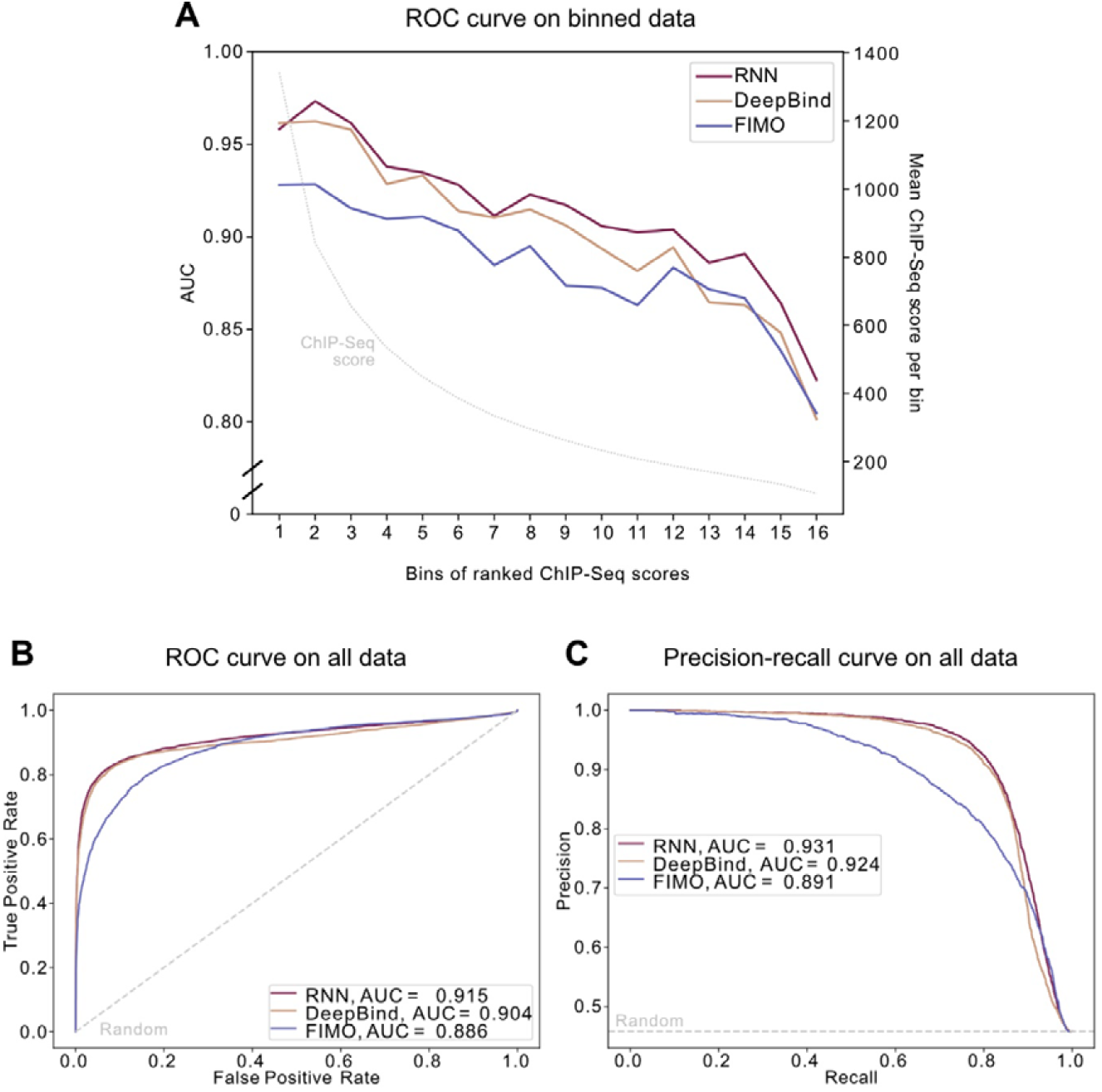
Performance of the RNN model on in vivo ChIP-Seq data. (A) Area under the curve (AUC) for the receiver operator characteristic (ROC) curve of the tested models on binned data. The 7,857 GRHL1 ChIP-Seq peaks were sorted by decreasing ChIP-Seq score and divided into bins of 500 sequences (357 in the last bin). 500 random MYOD peaks were selected as the negative dataset. The left y-axis corresponds to the AUC for each bin; the right y-axis and grey dotted line show the mean ChIP-Seq score per bin. AUCs are depicted as mean. (B) AUC for the ROC curve of the tested models on all 7,857 GRHL1 ChIP-Seq peaks. (C) AUC for the precision-recall curve of the tested models on all 7,857 GRHL1 ChIP-Seq peaks. The grey dotted lines in (B) and (C) depict the performance of a random guesser.

All models could discriminate between true and pseudo binding sites with an AUC higher than 0.75. Both deep learning approaches, the RNN and DeepBind, outperformed FIMO across nearly all bins, and they performed nearly equally for bins representing high ChIP-Seq scores. However, compared to our RNN, the performance of DeepBind dropped faster for bins representing lower ChIP-Seq scores that presumably reflect lower affinity binding sites (**Figure 4A**).

The RNN and DeepBind also showed the best performance when considering the whole dataset as determined by the AUC on the corresponding ROC (**Figure 4B**) and precision-recall curve (**Figure 4C**).

### The RNN model identifies novel GRHL1 binding sites

We hypothesised that high-scoring predictions of the RNN without predicted binding by FIMO could be used to identify previously unknown GRHL1 binding sites, which escape the more traditional PWM-based approach. To filter for such non-canonical binding sites identified by the RNN, we selected DNA sequences from the human GRHL1 ChIP-Seq dataset, which were clearly classified as binding by the RNN (RNN score > 0.99), and non-binding by FIMO (FIMO score < 0). These filtering steps identified 46 GRHL1 TFBSs identified exclusively by the deep learning-based RNN within the GRHL1 ChIP-Seq dataset (**Supplementary Table 5**).

### The RNN can accurately predict GRHL1 binding affinity

We selected one novel binding site identified by the RNN, from here on referred to as “Novel”, for further validation. For this sequence, the RNN predicted binding with very high confidence. Although DeepBind also identified a potential hit within the corresponding GRHL1 ChIP-Seq peak, the predicted binding sequence had a relatively low prediction score and differed from the sequence identified by the RNN (arrow heads in **Figure 3**).

The “Novel” sequence identified by the RNN completely lacked the highly conserved GRHL1 CNNG core motif, which so far has been considered obligatory for GRHL1-DNA binding (**Figure 5**) (28).

**Figure 5.**
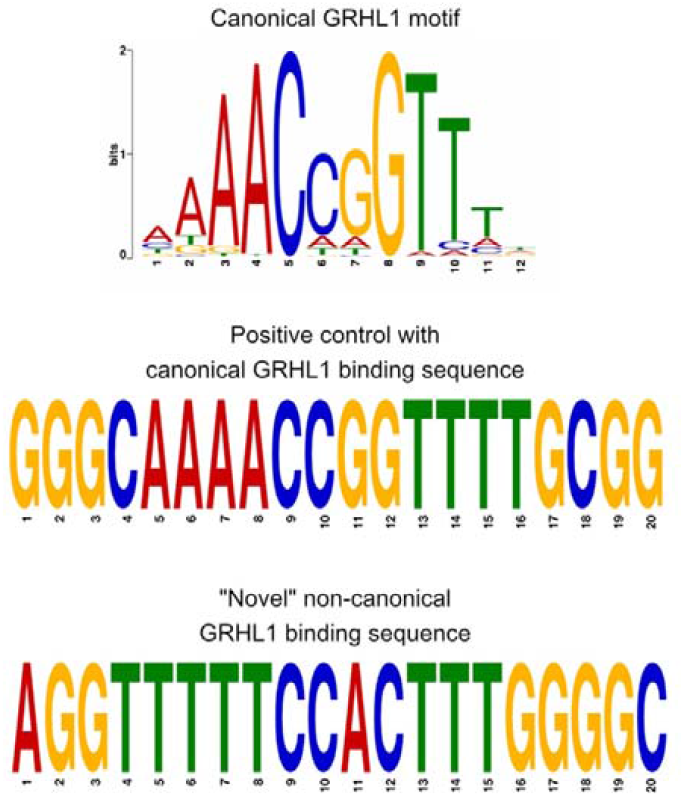
Comparison of the canonical GRHL1 motif (MA0647.1 from JASPAR), the positive control sequence containing the GRHL1 core motif, and the “Novel” non-canonical GRHL1 binding sequence lacking the CNNG core motif.

Microcalorimetric measurements of the binding affinity of GRHL1 towards this predicted non-canonical binding sequence determined a dissociation constant (K_d_) in the nanomolar range, indicating strong binding of substrate DNA (**Table 1, Supplementary Figure S1A**). A positive control (“Pos_Ctrl”), which is closely resembling the known GRHL1 binding motif AAACCGGTTT and for which both the RNN and FIMO predicted strong binding, showed a similar K_d_ as the “Novel” sequence (**Table 1, Supplementary Figure S2A**). For the negative control (“Neg_Ctrl”), selected from the non-binding MYOD dataset, no binding was observed (**Table 1**).

**Table 1.**
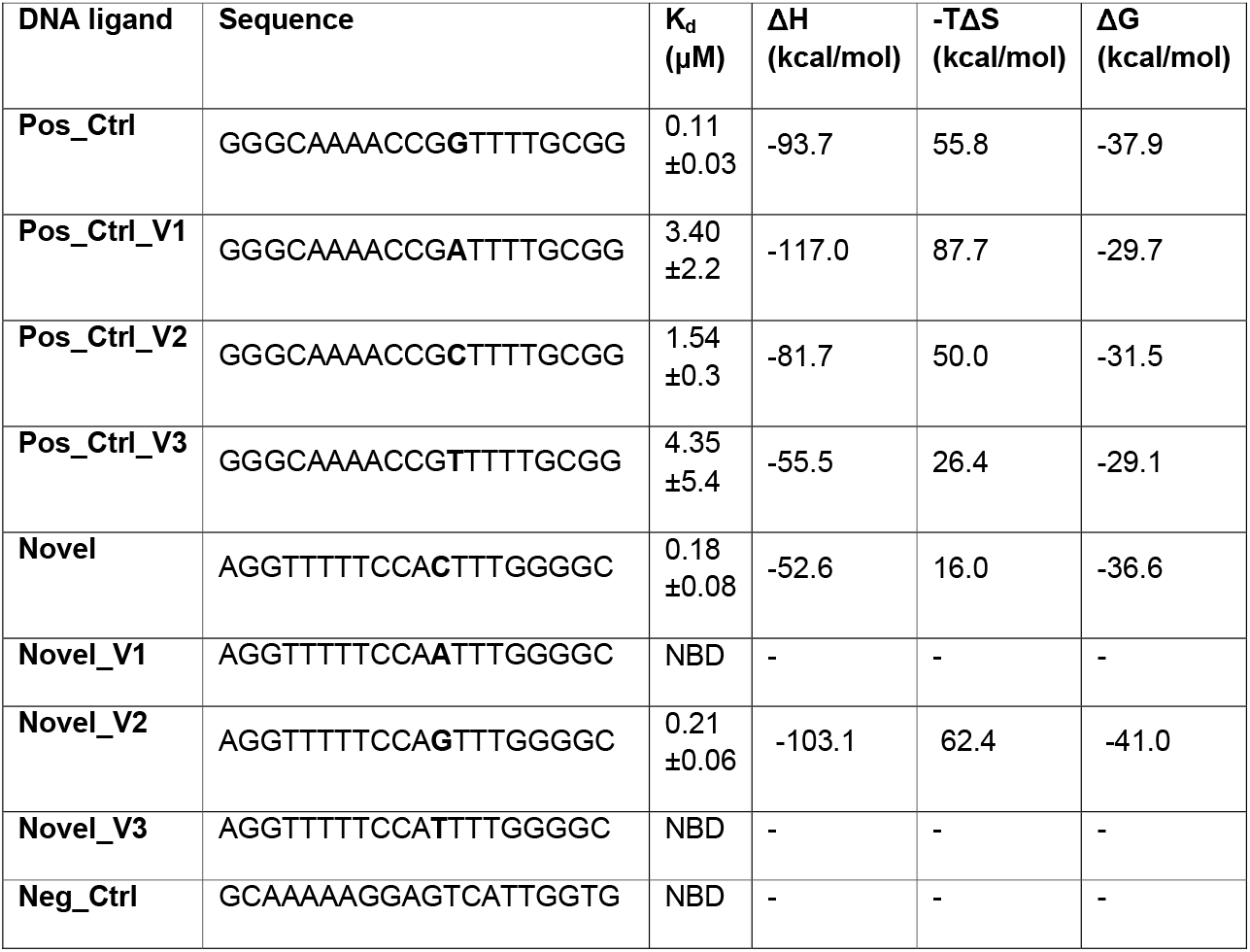
Results of isothermal titration calorimetry (ITC) experiment for the dsDNA sequences bound to GRHL1. The positive control (Pos_Ctrl) consists of the previously known GRHL1 binding motif. A MYOD sequence was used as a negative control (Neg_Ctrl). The position of the base variant is in bold. All characterised binding events are exothermic, and the Gibbs free energy of interaction is made by enthalpic contributions dominating over entropic ones. The binding affinities of variant sequences Pos_Ctrl_V1, Pos_Ctrl_V2, and Pos_Ctrl_V3 were lower by at least one order of magnitude compared to the Pos_Ctrl. For the Neg_Ctrl, Novel_V1, and Novel_V3 no binding was detected (NBD). K_d_ – dissociation constant; ΔH - enthalpy; -TΔS - entropy; ΔG - Gibbs free energy.

### The RNN can predict affinity changes caused by single nucleotide variants

We introduced all possible base exchanges into the positive control and “Novel” binding site and calculated the respective RNN binding scores to evaluate the RNNs’ ability to determine binding strength changes for single nucleotide variants (SNVs) within a binding site (**Supplementary Table 6**). We selected sequences with the mutation in position 12 for experimental validation based on the large differences in predicted binding strength between the original and the variant sequences (**Supplementary Table 7**). Obtained results proved binding of GRHL1 to high-scoring sequence variants, while sequences with an RNN score lower than 0.5 were below the experimental threshold for GRHL1 binding in our experiment (**Table 1, Supplementary Figure S1B-D**, and **2B-D**).

### RNN predictions highly correlate with experimentally determined GRHL1 binding affinity

For some experimentally tested sequences, we observed binding affinities lower by at least one order of magnitude compared to the positive control (**Table 1**). These sequences were considered binding by the RNN but showed a lower score. To estimate how accurately our RNN model can predict actual binding strength, we compared the predicted binding probability of the selected sequences of length 20 against the measured K_d_ values. Calculated correlation coefficients showed a strong correlation with a Pearson correlation of 0.926 (r2 = 0.858) and a corresponding Spearman correlation of 0.966 (r2 = 0.934) (**Figure 6A, Supplementary Table 8**). The correlation was much lower for DeepBind (**Figure 6B**) and FIMO (**Figure 6C**). This indicates that our RNN could predict the actual binding strength most closely and that RNN scores might serve as an indicator for the actual binding affinity (**Figure 6D, Supplementary Table 8**).

**Figure 6.**
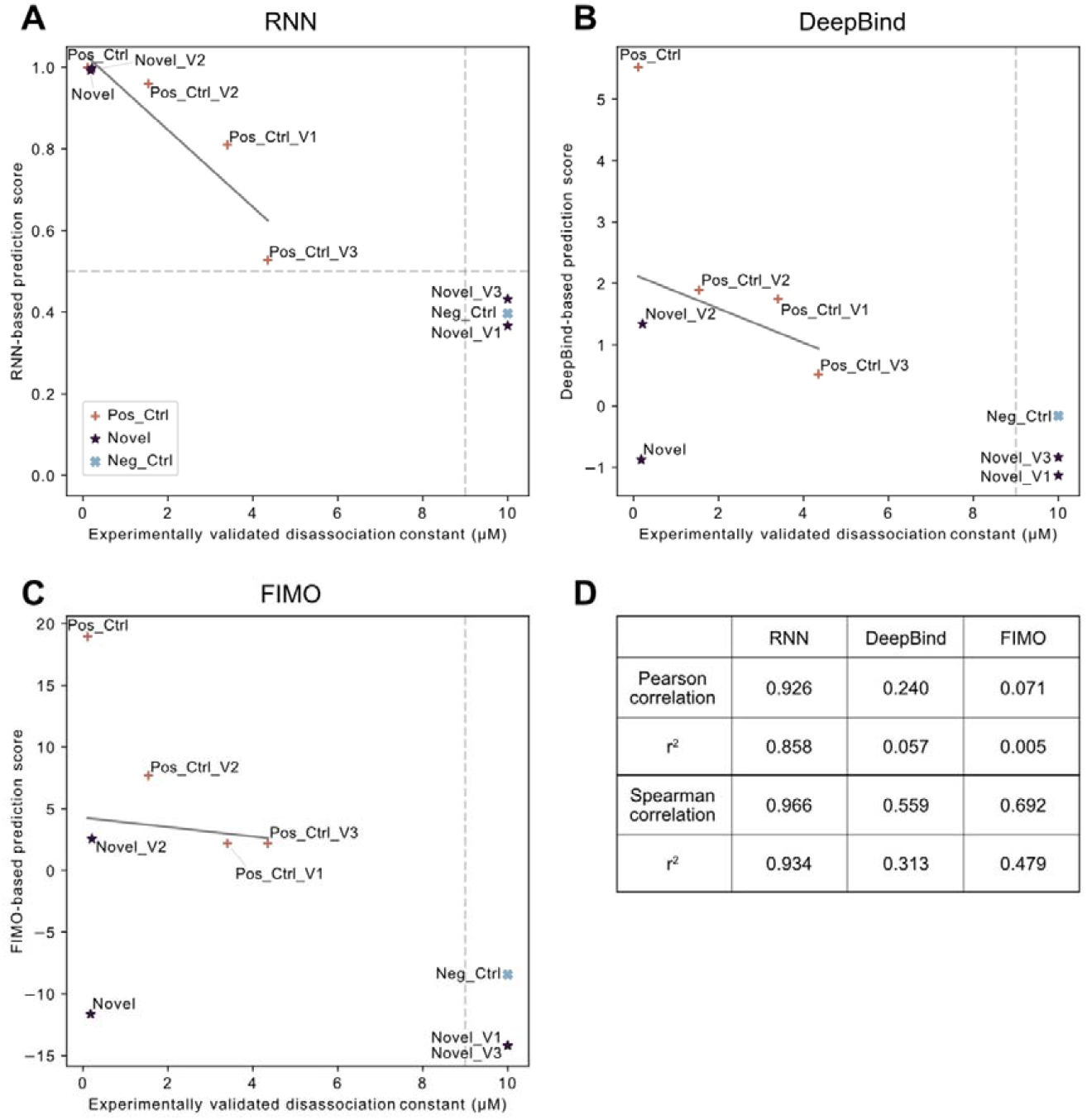
Correlation of experimentally determined dissociation constant (K_d_) and predicted binding scores: (A) RNN predictions. (B) DeepBind predictions. (C) FIMO predictions. (D) Correlation of prediction scores and K_d_ for all three models. A linear regression (grey line in plots) was used to determine the correlation. Pearson correlations calculated on raw values did not include non-binding sequences, while Spearman correlations calculated on ranked data considered all values. Non-binding sequences were set to a value of 10, as 9 is the detection threshold for the K_d_. Thresholds of binding versus non-binding are depicted as grey dotted lines; symbols represent the corresponding group.

## DISCUSSION

Transcription factors from the Grh family have been our focus for some time now. They play a significant role in embryonic development as well as tumour formation and progression (41,42). Thus, an in-depth understanding of their regulatory properties, including the identification of TFBSs throughout the genome, is crucial.

In this study, we trained a neural network using a combination of CNN and RNN architecture specifically for the transcription factor GRHL1, a representative of the Grh family. Our goal was to identify potential non-canonical TFBSs not represented by the known GRHL1 binding motif and to validate our predictions experimentally.

The ability of ANNs to learn longer-range interactions and interdependencies of all bases to one another adds a fundamental advantage over the traditional PWM approach in learning the syntax of TFBSs (14). PWMs are based on simple mathematical concepts of observed occurrences of nucleotides at certain positions within the sequence and are therefore easy to interpret. One of the greatest disadvantages of PWMs is their complete ignorance of interdependencies of the nucleotides within a sequence. PWMs tend to overrepresent TFBSs containing canonical sequences with high binding affinity and can have difficulties to accurately capture complex binding motifs (8). TFFMs and ANNs reduce the potential of missing functionally relevant low-affinity binding sites within the genome (9,43).

We compared the performance of our RNN to established methods based either on a PWM (FIMO) or a CNN model (DeepBind). Unsurprisingly, both our RNN and DeepBind performed generally better than the PWM-approach on both the HT-SELEX and the ChIP-Seq dataset. Although trained on the same data, our RNN performed slightly better than DeepBind and identified a new binding site not discovered by DeepBind. This improvement may be explained by the additional use of a recurrent layer in our model, which has been shown previously to perform better than a CNN alone (15-17). The recurrent layer allows the model to treat the bases of the binding sequences as ordered data, such that each position of the sequence is read in order. This enables the network to decide how much of the previous information should be retained during each subsequent step. This method is usually used in natural language processing settings. However, as properties similar to grammar and syntax can also be found in DNA the serial properties of RNNs can be advantageous in the context of TFBS identification within sequences (44-46).

It has been demonstrated before that ANNs can be utilized for identifying novel binding motifs or predicting the consequence of variants within a TFBS (14). Indeed, our RNN was able to identify 46 additional GRHL1 binding sites within a ChIP-Seq dataset that were missed entirely by FIMO. We experimentally confirmed a selected sequence as a novel GRHL1 binding site. Interestingly, this novel binding sequence shared very little similarity with the canonical GRHL1 motif AAACCGGTTT (MA0647.1 from JASPAR (7)) and lacked the CNNG core motif. The discovery of this alternative binding sequence in the current study seems particularly significant. Our recent biochemical and structural analysis of GRHL1 binding to double-stranded target DNA (28) fully explained the transcription factor’s high affinity for the canonical GRHL1 motif. The novel sequence was bound by GRHL1 with comparable affinity but must require an alternative protein-DNA interface with a new and unknown set of interatomic contacts.

In addition, our RNN was able to predict the effects of single nucleotide variants on GRHL1 binding affinity with higher accuracy than FIMO and DeepBind. RNN-based prediction scores highly correlated with experimentally determined GRHL1 binding affinities. Although the set of sequences for the correlation calculation was relatively small, the results indicate that RNN scores might serve as an indicator for the actual binding affinity and that our model is sensitive to single nucleotide variants.

Future studies of other transcription factors may benefit from this capability when exploring the role of disease-associated variants within hitherto unknown binding motifs.

## DATA AVAILABILITY

All code and scripts including all trained models are available in the GitLab repository: https://git-ext.charite.de/sebastian.proft/grhl1-binding

The motif MA0647.1 taken from JASPAR 2022 is available at: https://jaspar.genereg.net/matrix/MA0647.1/

HT-SELEX data used for training, validation, and testing can be found in the European Nucleotide Archive (ENA) under the project number PRJEB3289, SRA file ERR194761 (5): https://www.ebi.ac.uk/ena/browser/view/PRJEB3289

GRHL1 ChIP-Seq data can be found under the GEO accession number GSM4156315 and SRA accession number SRX7122002 (38).

MYOD ChIP-Seq data can be found under the GEO accession number GSM1218849 and SRA accession number SRX341010 (39).

## SUPPLEMENTARY DATA

Supplementary Data are available for download.

## ACKNOWLEDGEMENT

We would like to thank Minie Jung, who helped design the neural network.

## REFERENCES

1. Stormo, G.D. and Hartzell, G.W. (1989) Identifying protein-binding sites from unaligned DNA fragments. Proc. Natl. Acad. Sci., 86, 1183–1187.

2. Stormo, G.D. (2000) DNA binding sites: representation and discovery. Bioinformatics, 16, 16–23.

3. Tuerk, C. and Gold, L. (1990) Systematic Evolution of Ligands by Exponential Enrichment: RNA Ligands to Bacteriophage T4 DNA Polymerase. Science, 249, 505–510.

4. Johnson, D.S., Mortazavi, A., Myers, R.M. and Wold, B. (2007) Genome-Wide Mapping of in Vivo Protein-DNA Interactions. Science, 316, 1497.

5. Jolma, A., Yan, J., Whitington, T., Toivonen, J., Nitta, K.R., Rastas, P., Morgunova, E., Enge, M., Taipale, M., Wei, G. et al.. (2013) DNA-binding specificities of human transcription factors. Cell, 152, 327–339.

6. Badis, G., Berger, M.F., Philippakis, A.A., Talukder, S., Gehrke, A.R., Jaeger, S. A., Chan, E.T., Metzler, G., Vedenko, A., Chen, X. et al.. (2009) Diversity and Complexity in DNA Recognition by Transcription Factors. Science, 324, 1720–1723.

7. Castro-Mondragon, J.A., Riudavets-Puig, R., Rauluseviciute, I., Berhanu Lemma, R., Turchi, L., Blanc-Mathieu, R., Lucas, J., Boddie, P., Khan, A., Manosalva Pérez, N. et al.. (2022) JASPAR 2022: the 9th release of the open-access database of transcription factor binding profiles. Nucleic Acids Res., 50, D165–D173.

8. Siggers, T. and Gordân, R. (2014) Protein–DNA binding: complexities and multi-protein codes. Nucleic Acids Res., 42, 2099–2111.

9. Mathelier, A. and Wasserman, W.W. (2013) The Next Generation of Transcription Factor Binding Site Prediction. PLoS Comput. Biol., 9, e1003214.

10. Koo, P.K. and Ploenzke, M. (2020) Deep learning for inferring transcription factor binding sites. Curr Opin Syst Biol, 19, 16–23.

11. Zeng, Y., Gong, M., Lin, M., Gao, D. and Zhang, Y. (2020) A Review About Transcription Factor Binding Sites Prediction Based on Deep Learning. IEEE Access, 8, 219256–219274.

12. He, Y., Shen, Z., Zhang, Q., Wang, S. and Huang, D.S. (2021) A survey on deep learning in DNA/RNA motif mining. Brief Bioinform, 22.

13. Leiz, J., Rutkiewicz, M., Birchmeier, C., Heinemann, U. and Schmidt-Ott, K.M. (2021) Technologies for profiling the impact of genomic variants on transcription factor binding. Medizinische Genetik, 33, 147–155.

14. Alipanahi, B., Delong, A., Weirauch, M.T. and Frey, B.J. (2015) Predicting the sequence specificities of DNA- and RNA-binding proteins by deep learning. Nat Biotechnol, 33, 831–838.

15. Quang, D. and Xie, X. (2016) DanQ: a hybrid convolutional and recurrent deep neural network for quantifying the function of DNA sequences. Nucleic Acids Res., 44, e107–e107.

16. Pan, X., Rijnbeek, P., Yan, J. and Shen, H.-B. (2018) Prediction of RNA-protein sequence and structure binding preferences using deep convolutional and recurrent neural networks. BMC Genom., 19, 511.

17. Shen, Z., Bao, W. and Huang, D.S. (2018) Recurrent Neural Network for Predicting Transcription Factor Binding Sites. Sci Rep, 8, 15270.

18. Elman, J.L. (1990) Finding Structure in Time. Cognitive Science, 14, 179–211.

19. Nüsslein-Volhard, C., Wieschaus, E. and Kluding, H. (1984) Mutations affecting the pattern of the larval cuticle in Drosophila melanogaster. Wilhelm Roux’s Arch. Dev. Biol., 193, 267–282.

20. Bray, S.J. and Kafatos, F.C. (1991) Developmental function of Elf-1: an essential transcription factor during embryogenesis in Drosophila. Genes Dev, 5, 1672–1683.

21. Auden, A., Caddy, J., Wilanowski, T., Ting, S.B., Cunningham, J.M. and Jane, S.M. (2006) Spatial and temporal expression of the Grainyhead-like transcription factor family during murine development. Gene Expr Patterns, 6, 964–970.

22. Wilanowski, T., Caddy, J., Ting, S.B., Hislop, N.R., Cerruti, L., Auden, A., Zhao, L.-L., Asquith, S., Ellis, S., Sinclair, R. et al.. (2008) Perturbed desmosomal cadherin expression in grainy head-like 1-null mice. EMBO J, 27, 886–897.

23. Has, C. and Technau-Hafsi, K. (2016) Palmoplantar keratodermas: clinical and genetic aspects. J Dtsch Dermatol Ges, 14, 123–140.

24. Fabian, J., Lodrini, M., Oehme, I., Schier, M.C., Thole, T.M., Hielscher, T., Kopp-Schneider, A., Opitz, L., Capper, D., von Deimling, A. et al.. (2014) GRHL1 Acts as Tumor Suppressor in Neuroblastoma and Is Negatively Regulated by MYCN and HDAC3. Cancer Res, 74, 2604–2616.

25. Mlacki, M., Darido, C., Jane, S.M. and Wilanowski, T. (2014) Loss of Grainy Head-Like 1 Is Associated with Disruption of the Epidermal Barrier and Squamous Cell Carcinoma of the Skin. PLoS One, 9, e89247.

26. He, Y., Gan, M., Wang, Y., Huang, T., Wang, J., Han, T. and Yu, B. (2021) EGFR-ERK induced activation of GRHL1 promotes cell cycle progression by up-regulating cell cycle related genes in lung cancer. Cell Death Dis, 12, 430.

27. Nevil, M., Bondra, E.R., Schulz, K.N., Kaplan, T. and Harrison, M.M. (2017) Stable Binding of the Conserved Transcription Factor Grainy Head to its Target Genes Throughout Drosophila melanogaster Development. Genetics, 205, 605–620.

28. Ming, Q., Roske, Y., Schuetz, A., Walentin, K., Ibraimi, I., Schmidt-Ott, K.M. and Heinemann, U. (2018) Structural basis of gene regulation by the Grainyhead/CP2 transcription factor family. Nucleic Acids Res., 46, 2082–2095.

29. Whitfield, T.W., Wang, J., Collins, P.J., Partridge, E.C., Aldred, S.F., Trinklein, N.D., Myers, R.M. and Weng, Z. (2012) Functional analysis of transcription factor binding sites in human promoters. Genome Biol., 13, R50.

30. Weinhold, N., Jacobsen, A., Schultz, N., Sander, C. and Lee, W. (2014) Genome-wide analysis of noncoding regulatory mutations in cancer. Nat Genet, 46, 1160–1165.

31. Deplancke, B., Alpern, D. and Gardeux, V. (2016) The Genetics of Transcription Factor DNA Binding Variation. Cell, 166, 538–554.

32. Nishizaki, S.S., Ng, N., Dong, S., Porter, R.S., Morterud, C., Williams, C., Asman, C., Switzenberg, J.A., Boyle, A.P. and Hancock, J. (2019) Predicting the effects of SNPs on transcription factor binding affinity. Bioinformatics.

33. Grant, C.E., Bailey, T.L. and Noble, W.S. (2011) FIMO: scanning for occurrences of a given motif. Bioinformatics, 27, 1017–1018.

34. Scheich, C., Kümmel, D., Soumailakakis, D., Heinemann, U. and Büssow, K. (2007) Vectors for co-expression of an unrestricted number of proteins. Nucleic Acids Res., 35, e43–e43.

35. Gesell, T. and Washietl, S. (2008) Dinucleotide controlled null models for comparative RNA gene prediction. BMC Bioinformatics, 9, 248.

36. Bailey, T.L., Johnson, J., Grant, C.E. and Noble, W.S. (2015) The MEME Suite. Nucleic Acids Res., 43, W39–W49.

37. Budach, S. and Marsico, A. (2018) pysster: classification of biological sequences by learning sequence and structure motifs with convolutional neural networks. Bioinformatics, 34, 3035–3037.

38. Xu, G., Chhangawala, S., Cocco, E., Razavi, P., Cai, Y., Otto, J.E., Ferrando, L., Selenica, P., Ladewig, E., Chan, C. et al.. (2020) ARID1A determines luminal identity and therapeutic response in estrogen-receptor-positive breast cancer. Nat Genet, 52, 198–207.

39. MacQuarrie Kyle, L., Yao, Z., Fong Abraham, P., Diede Scott, J., Rudzinski Erin, R., Hawkins Douglas, S. and Tapscott Stephen, J. (2013) Comparison of Genome-Wide Binding of MyoD in Normal Human Myogenic Cells and Rhabdomyosarcomas Identifies Regional and Local Suppression of Promyogenic Transcription Factors. Mol. Cell. Biol., 33, 773–784.

40. Quinlan, A.R. and Hall, I.M. (2010) BEDTools: a flexible suite of utilities for comparing genomic features. Bioinformatics, 26, 841–842.

41. Kotarba, G., Taracha-Wisniewska, A. and Wilanowski, T. (2020) Grainyhead-like transcription factors in cancer – Focus on recent developments. Exp. Biol. Med., 245, 402–410.

42. Gasperoni, J.G., Fuller, J.N., Darido, C., Wilanowski, T. and Dworkin, S. (2022) Grainyhead-like (Grhl) Target Genes in Development and Cancer. Int. J. Mol. Sci., 23.

43. Trabelsi, A., Chaabane, M. and Ben-Hur, A. (2019) Comprehensive evaluation of deep learning architectures for prediction of DNA/RNA sequence binding specificities. Bioinformatics, 35, i269–i277.

44. Ji, S. (1999) The Linguistics of DNA: Words, Sentences, Grammar, Phonetics, and Semantics. Ann. N. Y. Acad. Sci, 870, 411–417.

45. Hie, B. D. Z.E., Berger, B. and Bryson, B. (2021) Learning the language of viral evolution and escape. Science, 371, 284–288.

46. Wahab, A., Tayara, H., Xuan, Z. and Chong, K.T. (2021) DNA sequences performs as natural language processing by exploiting deep learning algorithm for the identification of N4-methylcytosine. Sci. Rep., 11, 212.

