## Supplementary Figures for "Discovery of a non-canonical GRHL1 binding site using deep convolutional and recurrent neural networks"

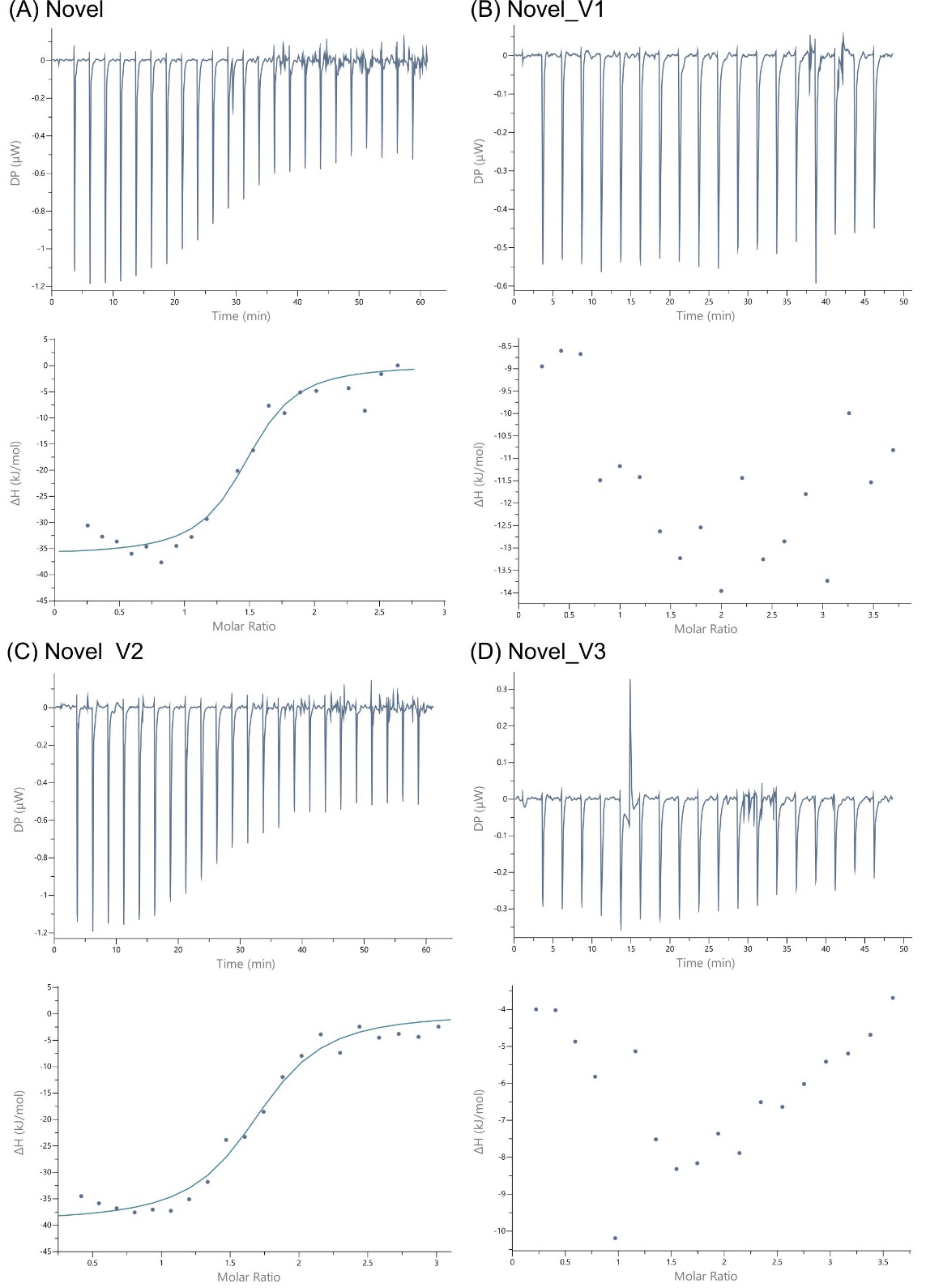

Figure S1: Calorimetric titration of Novel_V1-4 with GRHL1. The top panel shows the raw data obtained from 24 or 19 consecutive 1.5 or 2 μl injections of 165 μM or 200 μM GRHL1 solution into the sample cell containing ~10 μM of dsDNA. The binding isotherm of the bottom panel results from plotting heat peak areas against the dsDNA: protein molar ratio. The blue line represents the best fit of a model of one set of binding sites.

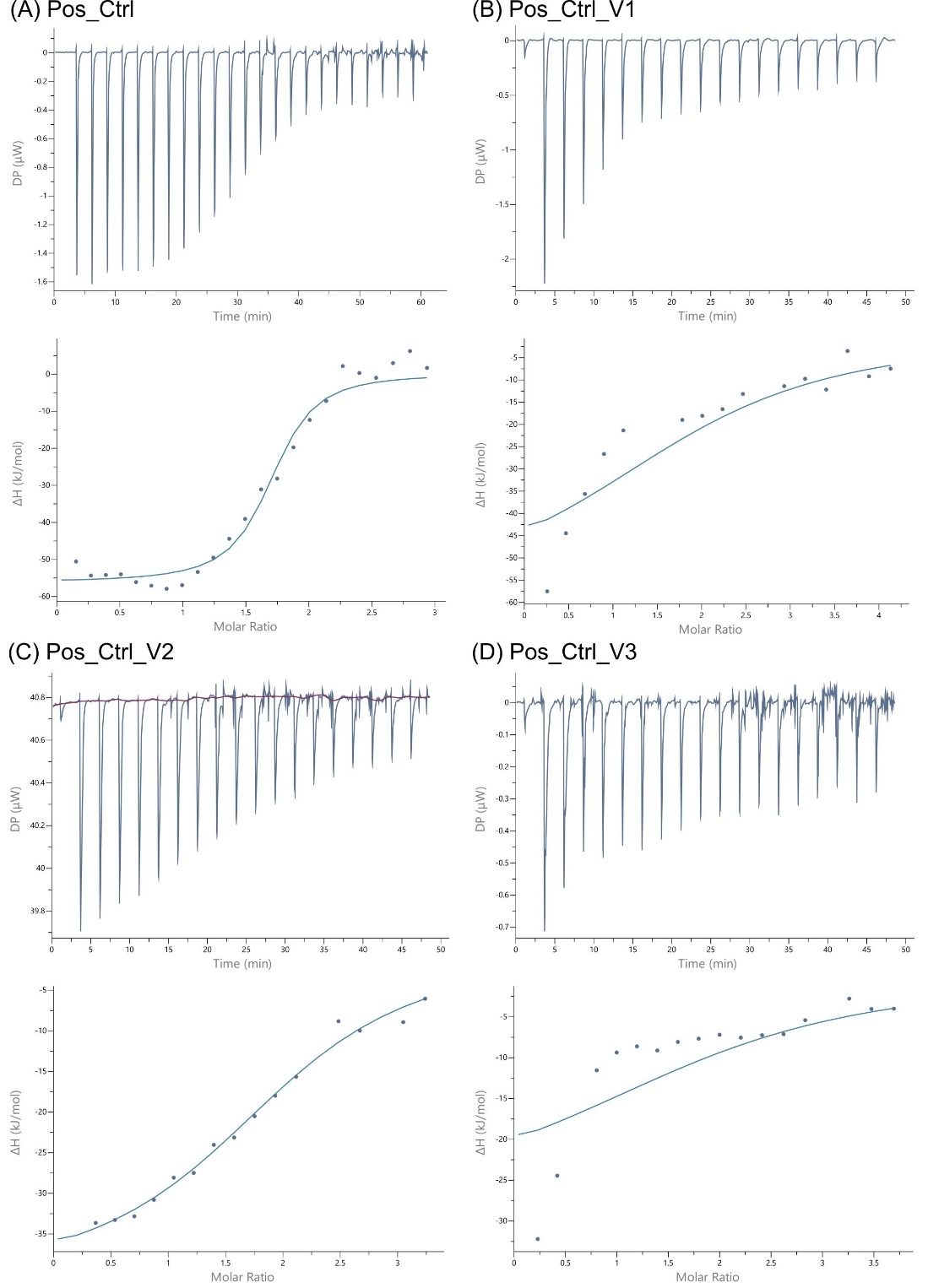

Figure S2: Calorimetric titration of Pos_Ctrl_V1-4 with GRHL1.The top panel shows the raw data obtained from 24 or 19 consecutive 1.5 or 2 μl injections of 165 μM or 200 μM GRHL1 solution into the sample cell containing ~10 μM of dsDNA. The binding isotherm of the bottom panel results from plotting heat peak areas against the dsDNA: protein molar ratio. The blue line represents the best fit of a model of one set of binding sites.
